# The genome of *Istocheta aldrichi* (Diptera: Tachinidae), a parasitoid of the Japanese beetle, *Popillia japonica* (Coleoptera: Scarabaeidae)

**DOI:** 10.1101/2025.11.02.686142

**Authors:** Pablo A. Stilwell, Jack A. Culotta, William D. Hutchison, Amelia R. I. Lindsey

## Abstract

*Istocheta aldrichi* Mesnil 1953 (Diptera: Tachinidae), is native to Japan, and has recently become an important biological control agent of the Japanese beetle, *Popillia japonica* (Coleoptera: Scarabaeidae), a pest with >300 host plants, including roses, linden trees, and numerous agricultural crops. During the past decade, *I. aldrichi*’s range has greatly expanded across North America, particularly in Quebec and Ontario, Canada, and in the Midwest U.S. In many areas, including Minnesota, 15-60% of Japanese beetles are parasitized by *I. aldrichi*, highlighting its importance as a natural enemy. To facilitate research on *I. aldrichi* and other tachinid flies we present a reference genome generated from a single individual. The final genome assembly is 875.3 Mbp contained in 1,041 scaffolds, with an N50 of 4.77 Mbp, and 99.5% complete Diptera BUSCOs present. We also present a complete mitogenome and use comparative genomics across 19 tachinid species to identify unique features of *I. aldrichi*.

Specifically, we find that while many tachinid lineages have experienced contractions in gene families, *I. aldrichi* is characterized by a relatively high number of gene family expansions, many of which are predicted to function in metal ion transport. Tachinids as a whole have undergone rapid copy number changes in 935 gene families, largely related to metabolism and morphogenesis. The *I. aldrichi* reference genome will further research opportunities on these parasitic flies, including their potential for biocontrol of *P. japonica*.

**ARTICLE SUMMARY:** The parasitic fly *Istocheta aldrichi* attacks and kills the Japanese beetle (*Popillia japonica*), a pest of more than 300 plants. There is potential to leverage *I. aldrichi* for biological control of the beetle, but application is hindered by a limited understanding of this fly’s biology. This reference genome for *I. aldrichi* will enhance research future efforts and our ability to manage *P. japonica*.

## INTRODUCTION

The parasitic fly, *Istocheta aldrichi* (Figure 1a), is a member of the family Tachinidae: one of the largest families in the order Diptera (true flies) and the largest family of parasitoids outside of Hymenoptera (Stireman III et al. 2006; Stireman III et al. 2019). As is characteristic of parasitoids, tachinids complete their development as parasites in or on a host but are free-living as adults. Compared to other tachinids with small eggs (microtype), *I. aldrichi* is known for its characteristic oviposition of large, macrotype, spherical white eggs placed on the pronotum (“back”) of the Japanese beetle (*Popillia japonica*, Coleoptera: Scarabaeidae) (Figure 1b)(Pelletier et al. 2023). Also known as the “Winsome fly”, the species is oviparous, endoparasitic, and is known for superparasitism, where up to eight eggs may be oviposited on a single beetle, especially females (Gagnon et al. 2023; Pelletier et al. 2023). The subfamily to which *I. aldrichi* belongs, Exoristinae, primarily consists of parasitoids of caterpillars (Stireman III et al. 2006). However, within the tribe Blondeliini, there have been a suite of host shifts to diverse and distantly related orders of hosts (Stireman III et al. 2019). As a specialist parasitoid of *P. japonica, I. aldrichi* is one such example. *Popillia japonica*, is an invasive pest that was first detected in the United States in 1916 and can damage over 300 plants (Shanovich et al. 2019; Clausen et al. 1927). Since invading, the beetle has spread throughout the eastern U.S. (Althoff and Rice 2022), with additional detections in several western states, and British Columbia, Canada – often prompting eradication efforts (Zhu et al. 2023; Makovetski and Abram 2024) Moreover, beginning in 2014, *P. japonica* was detected in Italy (Gotta et al. 2023; Pavesi 2014), with subsequent establishment in Switzerland (Graf et al. 2023). Early in the invasion process, eradication and crop protection relied heavily on pest trapping and insecticide use (Santoiemma et al. 2021; Venette and Hutchison 2021).

**Figure 1.**
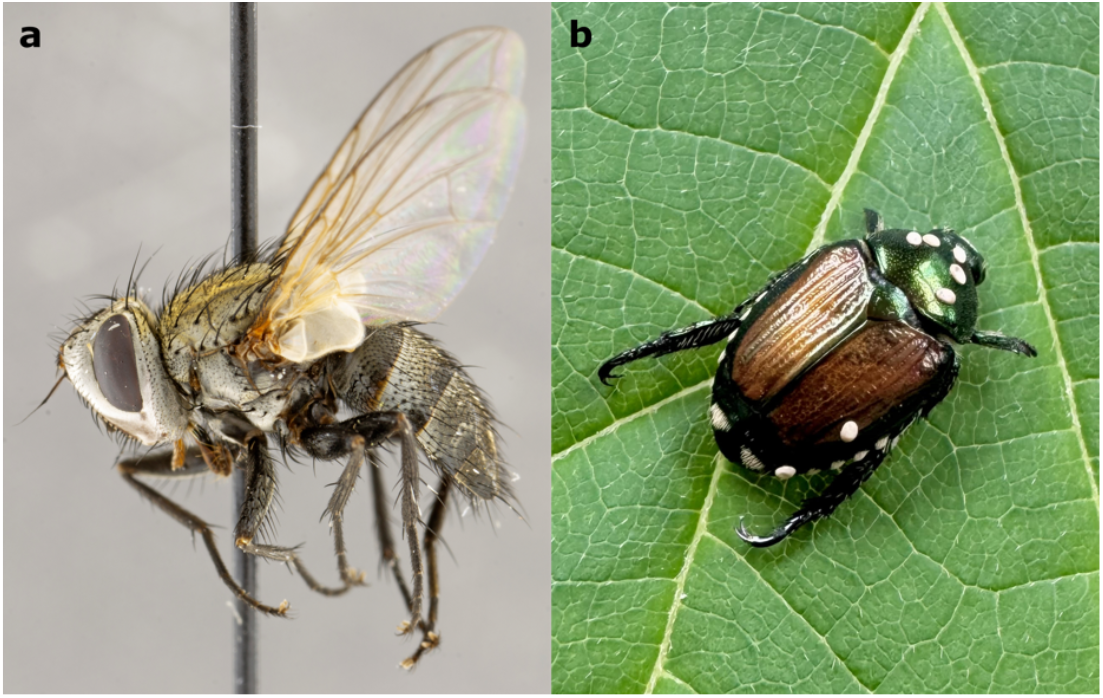
*Istocheta aldrichi*. **(a)** Female *I. aldrichi* (Canadian National Collection of Insects specimen ID CNC1711728). Image provided by the Canadian National Collection of Insects, Arachnids, and Nematodes (CNC), ©His Majesty The King in Right of Canada, as represented by the Minister of Agriculture and Agri-Food, licensed under the Open Government Licence Canada. **(b)** *I. aldrichi* eggs laid on the dorsum of its host, *Popillia japonica*. There are five eggs placed on the beetle’s pronotum (the typical oviposition location), and two additional eggs present on the right elytra (considered “misplaced”). Photo credit and permissions: Ellie R. Hutchison Cervantes.

However, for long-term sustainability, biological control, including the conservation or release of natural enemies, continues to be one of the most promising alternatives to chemicals for managing *P. japonica* (Althoff and Rice 2022; Abram et al. 2024; Brodeur et al. 2024).

As part of a classical biocontrol program in the 1920s, *I. aldrichi* was identified in Japan and released in several northeastern US states for *P. japonica* control (Clausen et al. 1927; Fleming 1968). Among the many parasitoid species released against *P. japonica, I. aldrichi* was deemed one of the most successful (Clausen et al. 1927; Fleming 1968). The recent range expansion of *I. aldrichi* in North America also portends considerable biocontrol potential (Gagnon et al. 2023; Makovetski and Abram 2024; Hutchison et al. 2024). *Istocheta aldrichi* exhibits several life history traits that are advantageous for biocontrol, including: (1) a tendency for high parasitism rates at low host densities, suggesting efficient searching behavior by females, (2) rapid mortality of host beetles following egg hatch of the fly (death within 5-7 d), and (3) that parasitized beetles stop feeding within 3-5 days of parasitization, which reduces defoliation and crop injury (Clausen et al. 1927; Brodeur et al. 2024; Hutchison et al. 2024).

Here, we provide a reference genome for *Istocheta aldrichi*: the first available for the tribe Blondeliini. As *I. aldrichi* continues to expand in range across North America, the parasitoid offers many biocontrol attributes that should facilitate a growing impact on one of the most damaging invasive pests, *P. japonica*. The genome sequence will provide a research base to assist with (1) evaluating biological characteristics, such as overwintering ability and diapause biology, (2) understanding the population genetics related to *I. aldrichi* establishment, (3) generating tools for accurate identification and monitoring, and (4) more broadly, improving our understanding of tachinid evolution.

## MATERIALS & METHODS

### Species Origin and Sampling Strategy

*Istocheta aldrichi* pupae were obtained from parasitized *P. japonica* beetles that had been field collected 14 days prior, from the foliage of wild grapes (*Vitis vinifera*), in Shoreview, MN, USA (45.1321 N, −93.11875 W), July 20, 2024, following published sampling procedures (Shanovich et al. 2021; Gagnon and Giroux 2019). This is the peak period of *I. aldrichi* activity in southern Minnesota (Hutchison et al. 2024), when the beetles are abundant, and beetles with eggs on their pronota are common. As noted by Brodeur et al. (2024), no other tachinid species to date, with similar oviposition behavior or egg shape, has been observed ovipositing on *P. japonica* adults in North America. Additionally, the high host specificity of *I. aldrichi* on *P. japonica* was recently supported (Makovetski et al. 2025). The species was also verified via the mitochondrial COX1 barcode (see below).

A cohort of 20 beetles, all having at least one characteristic *I. aldrichi* egg deposited on their pronota (Figure 1b), were placed individually in 475 ml plastic, ventilated cups, at 22°C and 16:8 (light:dark). Rearing cups were provisioned with fresh grape leaves for beetle nutrition (leaves changed daily), and moist filter paper for humidity. Although superparasitism (i.e., multiple parasitoid eggs deposited in or on a single host) by *I. aldrichi* is common, for the vast majority of cases, only one parasitoid larva successfully develops within the host’s thorax/abdomen (Clausen et al. 1927; Pelletier et al. 2023). At 14-20 days post-collection, the *I. aldrichi* larvae had either “cut out” of the host cadaver to pupate, or they had pupated inside the host. Within 24 hours after pupation, a total of six *I. aldrichi* pupae were collected and transferred to a −80°C freezer prior to DNA extraction. A single random pupa of unknown sex was selected for sequencing.

### Sequencing Methods and Sample Preparation

We extracted DNA from a single pupa using the Qiagen MagAttract kit following the manufacturer’s protocol. DNA was concentrated to 25 uL using Sergi Lab Supplies magnetic beads and the PacBio SRE kit was used to deplete fragments shorter than 10 kb. The sample was barcoded, library prepped with the ONT SQK-NBD114.24 kit, and sequencing was performed on an Oxford Nanopore P2 Solo instrument on a single flowcell (v.10.4.1). Libraries were recovered and flowcells were flushed with nuclease (EXP-WSH004 kit) prior to reloading every 24 hours. Data were basecalled using dorado v.0.7.3 and basecalling model dna_r10.4.1_e8.2_400bps_sup@v5.0.0.

### Nuclear Genome Assembly, Curation, and Quality Control

Reads were further processed with ‘dorado correct’ V.0.8.3+98456f7 and used for generating a primary assembly with Hifiasm v.0.19.9 (Cheng et al. 2021). The primary assembly was scaffolded using three rounds of ntLink v.1.3.11 (Coombe et al. 2023) with gap filling. Contamination screening and removal was performed using both NCBI Foreign Contamination Screens: (1) FCS-adaptor to remove adaptor contamination, and (2) the FCS-GX Genome Cross-Species Aligner (Astashyn et al. 2024) with the taxonomic ID NCBI:txid2500616 (*Istocheta aldrichi*). Purge_dups v1.2.5 was used to remove redundant haplotigs and overlaps from the genome assembly (Guan et al. 2020). Genome completeness was assessed with Compleasm v0.2.6 (Huang and Li 2023), which scored assemblies against the Diptera_odb10 database of 3,285 benchmarking single-copy orthologs (BUSCOs) and Merqury v1.3 (Rhie et al. 2020).

### Repeat Assembly Techniques

Repeat families were identified *de novo* using RepeatModeler v2.0.1 (Flynn et al. 2020). RepeatMasker v4.1.1 (Tarailo-Graovac and Chen 2009) was used to soft mask genomes with the *de novo* generated repeat libraries using slow search mode.

### Gene Finding Methods

Gene prediction was performed with BRAKER v3.0.8 (Brůna et al. 2021) using soft masked genomes against a curated protein database of Arthropoda proteins from OrthoDB (Kuznetsov et al. 2023). Gene function and domain annotation was performed by InterProScan v5.75-106.0 (Jones et al. 2014) and eggNOG-mapper v2.1.13 (Huerta-Cepas et al. 2019), utilizing the following databases: AntiFam v8.0, CDD v3.21, Coils v2.2.1, FunFam v4.3.0, Gene3D v4.3.0, Hamap v2025_01, MobiDBLite v4.0, NCBIfam v17.0, PANTHER v19.0, Pfam v37.4, PIRSF v3.10, PIRSR v2025_01, PRINTS v42.0, ProSitePatterns v2025_01, ProSiteProfiles v2025_01, SFLD v4, SMART v9.0, SUPERFAMILY v1.75, and eggNOG v5.0.2. Functional annotations and database references including Gene Ontology (GO) terms from the two programs were merged with the structural gene annotation from BRAKER to produce the final generic feature file.

### Comparative Genomics and Gene Family Evolution

Genomes of 18 other tachinids plus two outgroups (from the families Polleniidae and Calliphoridae) were used in comparative analyses (Table 1). Genome assemblies were retrieved from NCBI and each assembly was annotated following the same pipeline as described above for *I. aldrichi*, including *de novo* repeat identification and masking, and annotation with BRAKER. Orthologous groups of proteins (i.e., gene families) were clustered with OrthoFinder v2.5.4 (Emms and Kelly 2019), based on the longest transcript variant for each protein coding gene. The species tree generated by OrthoFinder was converted to an ultrametric tree via the chronos function in the ape package v.5.8-1 (Paradis et al. 2019), and used in combination with the gene family counts to estimate significant gene family contractions and expansions across the phylogeny with CAFE v5.1 (Mendes et al. 2020). CafePlotter v0.2.0 (https://github.com/moshi4/CafePlotter) was used to extract gene families from the resulting

**Table 1.**
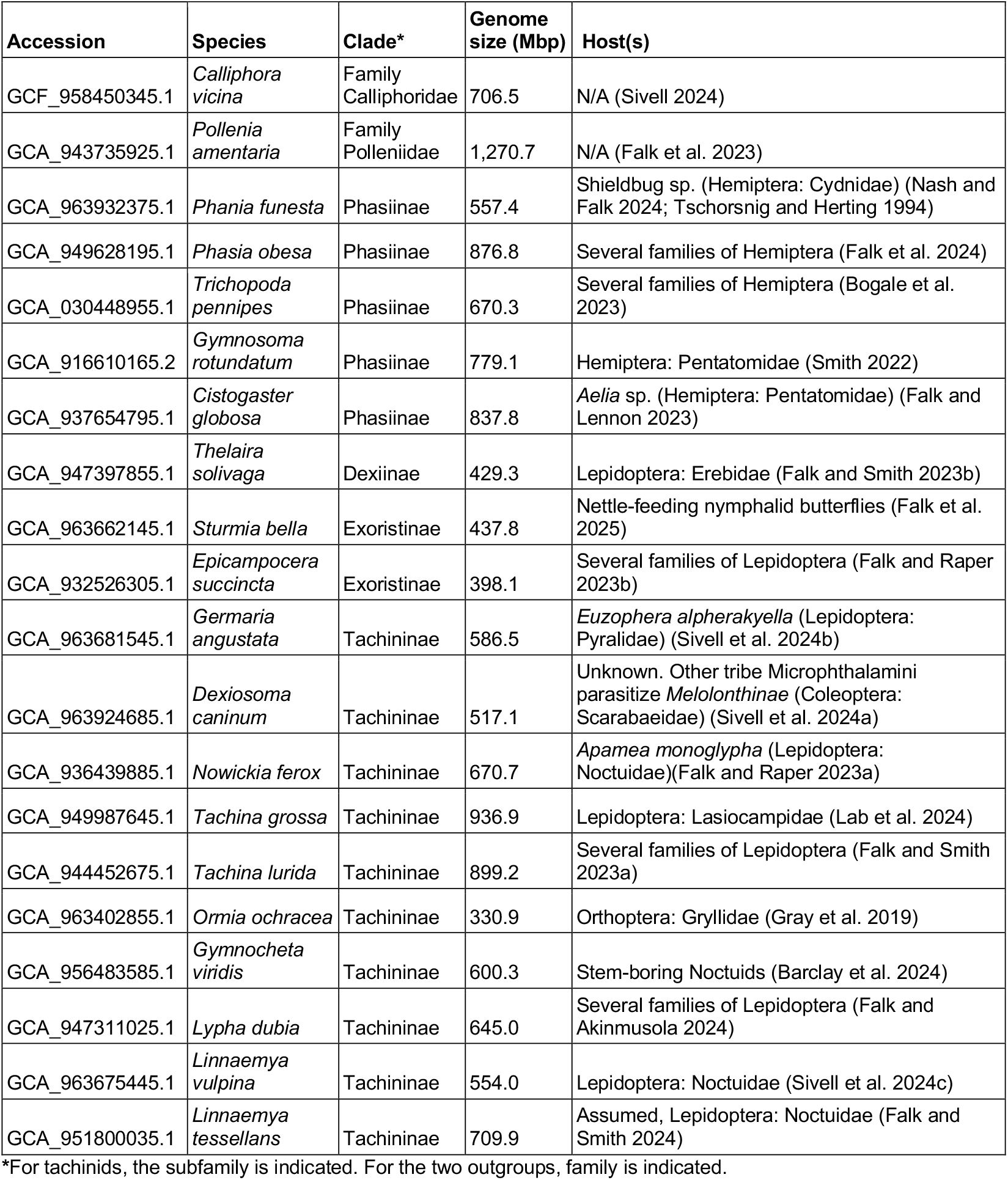
Genomes used in comparative analyses.

CAFE output files. To determine significant enrichments of gene ontology (GO) terms within sets of genes or gene families, we followed previously developed methods (Lindsey et al. 2018). In brief, statistical testing was performed with BiNGO v 3.0.5 (Maere et al. 2005), implemented in Cytoscape v3.9.1 (Shannon et al. 2003), using hypergeometric tests and Benjamini & Hochberg FDR correction, at a corrected significance level of 0.05. The background set of GO terms differed depending on the comparison. For comparisons within one genome, the background included all genes in that genome. To test for GO term enrichment in a set of gene families, we first annotated all proteins used in orthogroup clustering with OrthoFinder via the EMBL web eggNOG mapper-2.1.12 (http://eggnog-mapper.embl.de/) to generate GO terms.

Then, we created a custom background set of GO terms for each gene family, in which the GO terms included those represented by at least 40% of the genes in that family.

### Mitochondrial Genome

The mitochondrial genome was assembled using MitoHiFi v3.2.2 (Uliano-Silva et al. 2023) using the dorado-corrected reads as input, and the findMitoReference function by specifying the species *Istocheta aldrichi* and min_length 14000. The MitoHifi assembled genome was circularized using Circlator v1.5.5 fixstart function (Hunt et al. 2015) and annotated using MITOS2 implemented on the Galaxy web server (Bernt et al. 2013; Donath et al. 2019). The resulting COX1 sequence was extracted and queried against the Barcode of Life Database (BOLD; https://boldsystems.org/) to further validate sample identity.

## RESULTS & DISCUSSION

### Sequencing and Assembly

We extracted high molecular weight DNA from a single *I. aldrichi* pupa and used Oxford Nanopore sequencing to generate a total of 10.6M reads totaling 49.9 Gbp (Supplemental Table S1). Error corrected reads were then used to assemble a draft genome with Hifiasm. The draft genome of 916.9 Mbp was contained in 2,297 contigs with an N50 of 3.18 Mbp (Table 2). After scaffolding, decontamination, and purging haplotigs, the final genome assembly was 875.3 Mbp, contained in 1,041 scaffolds (1,063 contigs), with a scaffold N50 of 4.77 Mbp (Table 2).

**Table 2.** *Istocheta aldrichi* genome assembly and curation statistics.

| Metric | Draft<sup>#</sup> | Final (contigs) | Final (scaffolds) |
| --- | --- | --- | --- |
| Sequences | 2,297 | 1,063 | 1,041 |
| Total assembly length (bp) | 916,858,228 | 875,160,985 | 875,256,404 |
| Min sequence length | 5,340 | 5,340 | 5,340 |
| Mean sequence length | 399,155 | 823,293 | 840,784 |
| Max sequence length | 16,220,652 | 17,929,400 | 17,929,400 |
| N50 | 3,182,884 | 4,766,515 | 4,769,057 |
| L50 | 81 | 56 | 55 |
| %GC | 30.62 | 30.42 | 30.42 |
| Compleasm* | C:99.5%, 3270<br>[S:97.96%, 3218<br>D:1.58%, 52]<br>F:0.06%, 2<br>I:0.00%, 0<br>M:0.40%, 13<br>N:3285 | C:99.5%, 3269<br>[S:98.33%, 3230<br>D:1.19%, 39]<br>F:0.09%, 3<br>I:0.00%, 0<br>M:0.40%, 13<br>N:3285 |  |

<sup>#</sup> Draft assemblies are from HiFiasm, before ntLink and before purging contaminants.
\* Standard BUSCO annotation: Complete dipteran BUSCOs (C) [Complete, single-copy BUSCOs (S), Complete, duplicated BUSCOs (D)], Fragmented BUSCOs (F), Interrupted BUSCOs (I) Missing BUSCOs (M), Total BUSCO groups searched (N).

The final genome assembly was relatively complete, with a 99.5% Compleasm score for dipteran BUSCOs. At 875.3 Mbp, *I. aldrichi* has the fourth largest genome of sequenced tachinid species (n=22). Compared to similarly sized tachinid genomes (e.g., *Phasia obesa*, 876.8 Mbp, see Table 1), this assembly of *I. aldrichi* has fewer contigs (1,063 versus 3,378) and a larger contig N50 (4.77 Mbp versus 0.47 Mbp, respectively). While *I. aldrichi* has not been scaffolded onto chromosomes like many of the other tachinid genomes (including *P. obesa*), the *I. aldrichi* assembly here is nevertheless of high quality.

### Mitochondrial Genome

We assembled and annotated a complete 18,469 bp mitochondrial genome from *I. aldrichi*. We identified a complete gene set including small and large rRNAs, 22 tRNAs, and 13 protein coding genes (Figure 2). The gene arrangement, like most other dipterans and tachinids, is the same as the ancestral insect mitochondrial genome (Cameron 2014; Pei et al. 2024). We cross-referenced the COX1 sequence from the mitochondrial genome assembly with the BOLD barcode database and determined that the mitogenome sequenced here was identical to published *I. aldrichi* barcodes at COX1 (BIN BOLD:ADE2384, see also Shanovich et al. (2021)).

**Figure 2.**
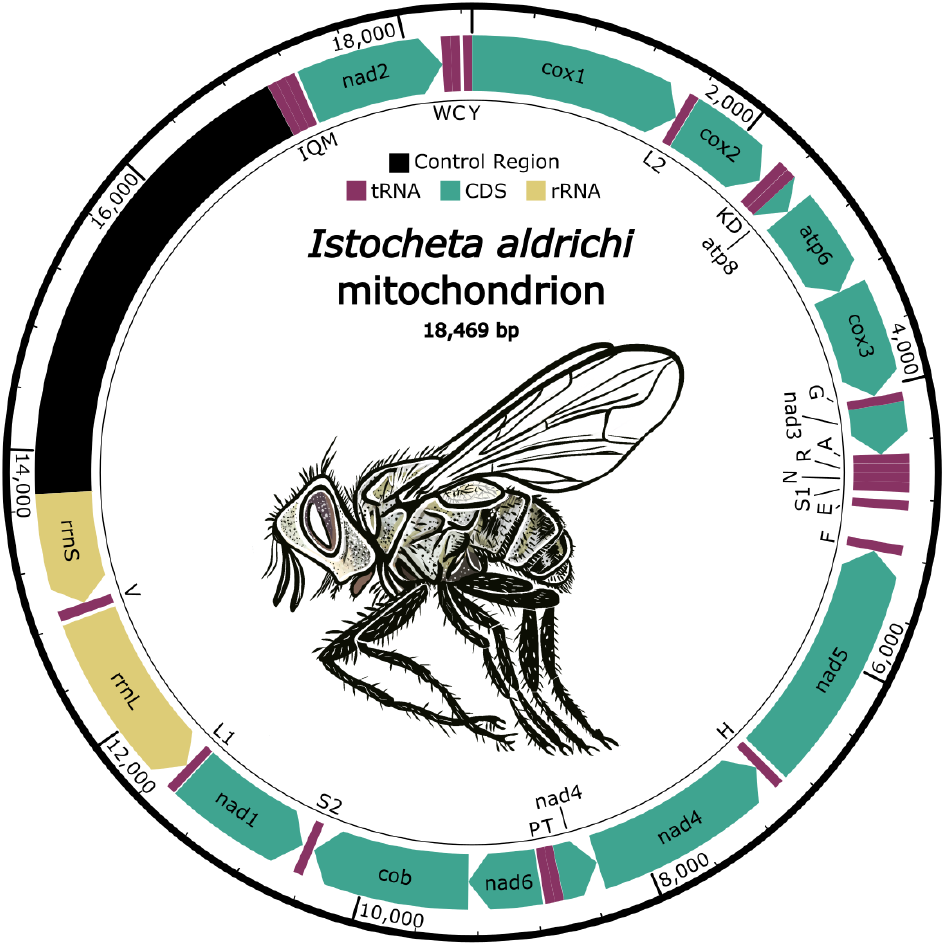
*Istocheta aldrichi* mitochondrial genome. (*I. aldrichi* drawing: Melissa Schreiner, Colorado State University-Extension, Grand Junction, CO). The complete mitochondrial genome of *I. aldrichi*, with annotated rRNAs, tRNAs, coding sequences (CDS), and the predicted control region (defined based on relatives (Cameron 2014; Pei et al. 2024). tRNAs are indicated with single-letter IUPAC-IUB abbreviations corresponding to the amino acid. The mitochondrial genome and annotations are available at NCBI under accession number PX213662.

### Repeats

Greater than 70% of the *I. aldrichi* genome was derived from repetitive elements (Table 3). The majority of repeats, corresponding to 50.6% of the genome length, were unclassified, which is not atypical for non-model insect species (Sproul et al. 2023; Petersen et al. 2019). Retroelements and DNA transposons were the largest categories of identified elements, and accounted for 8.4% and 10% of the genome, respectively.

**Table 3.** Repetitive sequences in *Istocheta aldrichi*.

| Name | Number | Length (bp) | Percent (%) |
| --- | --- | --- | --- |
| Retroelements | 126,476 | 73,636,889 | 8.41 |
| Penelope class | 7,988 | 3,512,242 | 0.40 |
| LINE class | 92,362 | 45,926,233 | 5.25 |
| L2/CR1/Rex | 28,620 | 10,768,381 | 1.23 |
| R1/LOA/Jockey | 4091 | 2,630,147 | 0.30 |
| R2/R4/NeSL | 95 | 102,292 | 0.01 |
| LTR class | 34,114 | 27,710,656 | 3.17 |
| BEL/Pao | 6,335 | 5,393,792 | 0.62 |
| Ty1/Copia | 1,751 | 1,533,512 | 0.18 |
| Gypsy/DIRS1 | 25,970 | 20,764,846 | 2.37 |
| DNA transposons | 197,900 | 87,342,017 | 9.98 |
| hobo-Activator | 57,782 | 17,658,477 | 2.02 |
| Tc1-IS630-Pogo | 73,612 | 33,272,189 | 3.80 |
| Rolling-circles | 171 | 42,189 | 0.00 |
| Unclassified | 1,900,031 | 442,956,545 | 50.61 |
| Total interspersed repeats |  | 603,935,451 | 69.01 |
| Simple repeats | 168,984 | 8,707,703 | 0.99 |
| Low complexity | 41,205 | 1,981,500 | 0.23 |
| Bases masked |  | 614,668,413 | 70.23 |

### Gene Finding

Structural annotation of the *I. aldrichi* genome characterized 32,005 mRNAs representing 28,575 genes (Table 4), which is only slightly above the average gene number for tachinids based on our annotations (mean = 25,816; Supplemental File S2). In contrast, the cricket parasitoid *Ormia ochrace* had the fewest genes (n*=*14,670) whereas *Cistogaster globosa*, a stinkbug specialist, was the most gene rich of the tachinids (n=34,717, Figure 3b). A total of 27,877 *I. aldrichi* proteins were functionally annotated by eggNOG-mapper, with 2,559 annotated by eggNOG-mapper alone. Using InterProScan, 27,692 proteins were functionally annotated, with 2,374 characterized by InterProScan but not eggNOG-mapper. Finally, 25,318 proteins were functionally annotated by both programs. Results from both functional annotation programs were merged.

**Table 4.** *Istocheta aldrichi* genome annotation metrics.

| Metric | Value |
| --- | --- |
| Number of genes | 28,575 |
| Number of mRNAs | 32,005 |
| Number of exons | 97,869 |
| Number of introns | 65,864 |
| Mean exons per mRNA | 3 |
| Total gene length | 162,793,239 bp |
| Longest gene | 224,090 bp |
| Mean gene length | 5,697 bp |
| Longest CDS | 66,945 bp |
| Mean CDS length | 1,280 bp |
| Longest exon | 32,355 bp |
| Mean exon length | 427 bp |

**Figure 3.**
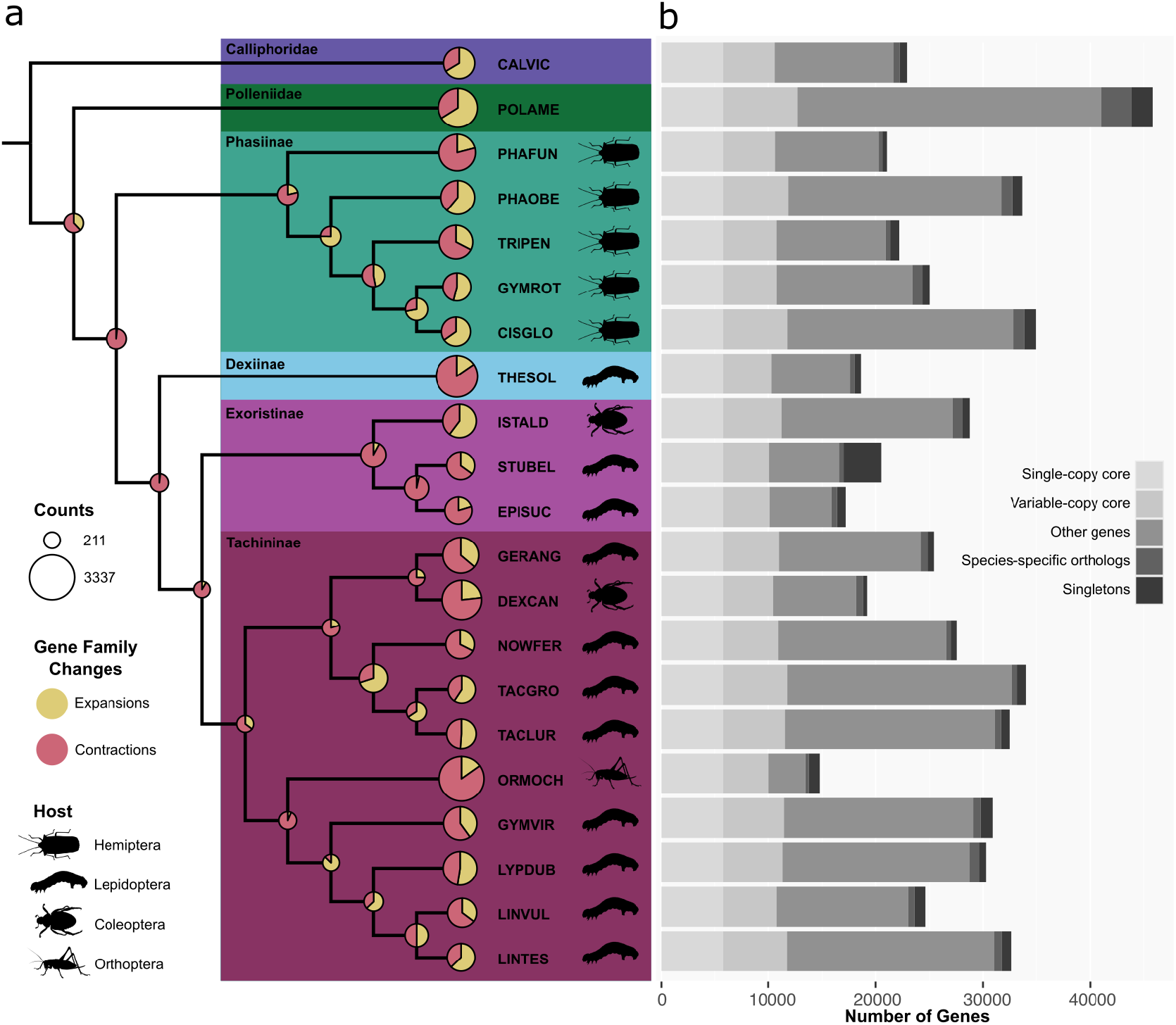
Tachinid genome evolution. **(a)** Phylogeny representing all Tachinidae subfamilies and select outgroups, showing superfamily wide gene family size changes and tachinid host associations. Species abbreviations correspond to the first three letters of the genus and first three letters of the species epithet. **(b)** Stacked bar-chart reflecting the composition of genes associated with different levels of gene family conservation.

### Comparative Genomics of Tachinids

We leveraged the published genomes of 18 additional tachinids and two outgroup species for comparative analyses (Table 1, Figure 3). Phylogenetic reconstruction recapitulated most relationships that are well-supported by more in-depth analyses of Diptera and Tachinidae evolution (Stireman III et al. 2019; de Paula et al. 2024). Specifically, each tachinid subfamily was recovered as monophyletic, and the sister relationships of the subfamilies agreed with recent published phylogenies, except for the placement of the single representative species of Dexiinae, as this subfamily is generally accepted to be sister to Phasiinae (Stireman III et al. 2019; de Paula et al. 2024). *Istocheta aldrichi* was sister to the other two species within the Exoristinae (*Sturmia bella* + *Epicampocera succincta*), and Exoristinae was sister to Tachininae (Figure 3). Across the 21 genomes, 538,711 genes (96.4% of the total 558,763) were clustered into 24,885 gene families (Supplemental Tables S2,S3). The largest gene family contained 1,305 genes, had between 1 and 208 paralogs per species, and was represented by all species except for *Sturmia bella*. 8,935 gene families contained genes from all species, and of these, 5,707 gene families consisted entirely of single-copy genes. 3,565 gene families (15,289 genes in total) contained genes exclusively derived from the same genome (i.e., species-specific families). 9,001 gene families were present in all tachinid species, only one of which was also not present in both outgroups. The subfamily to which *I. aldrichi* belongs, Exoristinae, were uniquely missing four gene families that were present in all other species. All Exoristinae species also had genes in 33 gene families that were unique to the Exoristinae. Finally, for *I. aldrichi*, 97.5% of the 28,575 genes were assigned to gene families, and of these, there were 878 genes in 211 species-specific gene families. 702 *Istocheta aldrichi* genes were defined as singletons, and there were 81 gene families that were uniquely lost in this lineage but present in all other species.

Curiously, one of the gene families missing in Exoristinae encoded for centromeric H3 (CENH3, “Cid” in *Drosophila melanogaster*, “CEN-P” in yeast). On further inspection we determined that the Exoristinae CENH3 proteins were not missing, but instead clustered with canonical H3 variants of the other species due to their short N-terminus relative to the other tachinid CENH3 proteins. These Exoristinae proteins were clearly identifiable as centromeric H3 variants due to the Q69A and F85Y amino acid substitutions (relative to the highly conserved canonical H3), and a longer loop 1 region in the histone fold domain by one amino acid (Drinnenberg et al. 2014; Malik and Henikoff 2003). However, there did not appear to be any Exoristinae canonical H3 proteins present in that same gene family. Blastp searches against the annotated *I. aldrichi* proteome (query: *Drosophila melanogaster* H3, GenBank CAA32434.1) similarly did not recover any canonical H3 proteins, only CENH3. Finally, we determined that the H3 proteins were encoded in the genome, but they had undergone numerous duplications to the point that they had been masked during repetitive element analyses. Indeed, we identified 131 open reading frames across five contigs which encoded for identical H3 proteins. While having numerous gene copies of histones is not unusual (Rooney et al. 2002), it seems the recent copy number expansion in this lineage is distinct from other clades in the family and also paired with a more divergent CENH3 N-terminus. Perhaps related to the signatures of histone evolution, the singleton genes unique to *I. aldrichi* were significantly enriched for eight GO terms, six of which were related to mitosis and DNA repair (Supplemental Table S3) such as G2/M cell cycle checkpoints and double-stranded break repair and processing. In contrast to the *I. aldrichi* singletons, genes in *I. aldrichi*-specific gene families were significantly enriched for metabolic functions, especially those related to vitamin A, terpenoids, protein catabolism, mitochondrial biology, and the circulatory system (Supplemental Table S4).

### Gene Family Size Evolution

In addition to identifying gene families that had been completely gained or lost, we also identified gene families that underwent changes in copy number at specific points across the phylogeny and characterized each one as either “expanding” (e.g., gaining paralogs) or “contracting” (e.g. losing paralogs) at a given node or leaf (Figure 3a). First, we identified 935 gene families which experienced significant changes in the rate of gene gain and loss across the tachinid phylogeny. These families were significantly overrepresented for a suite of GO terms, largely involving metabolic functions and morphogenesis (Supplemental Table S5), which may relate to the evolution of parasitism in this group, especially in the context of host switches. We determined that at the root of Tachinidae, 618 gene families changed in size, and these were primarily contracting families (n=607). In fact, this node had the lowest percentage of expanding gene families (relative to all changing families at a node) across the phylogeny. The nodes representing the ancestors of each subfamily were also characterized by a relatively high level of gene family contractions (C) relative to expansions (E): Phasiinae [C:516, E:140], Dexiinae [C:2,535, E:462], Exoristinae [C:1,074, E:92], Tachininae [C:169, E:92]. In contrast, *I. aldrichi* had an especially high number of gene families that expanded in size on this branch (n=1,261); the second highest number across the extant species (second to *Phasia obesa*, n=1,302). An additional 833 gene families were determined to have contracted in the *I. aldrichi* lineage. *Istocheta aldrichi* was one of only four tachinid species where >60% of significant families underwent lineage-specific expansions (along with *Phasia obesa, Linnaemya tessellans*, and *Cistogaster globosa*). The gene families that increased in size on the *I. aldrichi* branch were significantly overrepresented for five GO terms (Supplemental Table S6), four of which were related to copper/metal ion transport. The fifth overrepresented GO term was for RNA dependent DNA replication, likely indicative of viral genes. Finally, gene families that contracted in size on the *I. aldrichi* branch were significantly overrepresented for 17 GO terms, all of which related to neurological processes such as sensory perception, mating/reproductive behavior, and cognition (Supplemental Table S7).

### SUMMARY

The sequencing and analyses of the *I. aldrichi* genome presented herein represent the first available reference genome for the tribe Blondeliini, and the first functional genomic comparisons across the family Tachinidae, the second largest dipteran family in terms of numbers of species descriptions (Stireman III et al. 2019). For *I. aldrichi*, this will facilitate further research on the biological control of *P. japonica*. Of particular interest, this reference genome will support research on population genetics related to the recent spread of this fly, and mechanistic investigations of its ability to target *P. japonica*, even at low densities. In addition to the importance of tachinids as biological control agents of many insect pests (Grenier 1988; Rezaei et al. 2022), their evolutionary history affords myriad opportunities to understand rapid speciation, parasitism, and major transitions between feeding ecologies and host associations (Stireman III et al. 2019). Indeed, others have estimated that Tachinidae may be one of the most rapidly diversifying lineages across all of metazoa (Scholl and Wiens 2016), and our work here provides a foundation to explore this diversity.

## Data Availability

BioProject: PRJNA1297203

BioSample: SAMN50213581

Sequencing Reads: SRR34721792

Genome: #forthcoming, submitted to NCB

I Mitochondrial genome: PX213662

Supplemental File S1. Rmarkdown file containing all code

Supplemental File S2. gzip file containing structural annotations (gff3) for all species (via figshare)

Supplemental Table S1. Sequencing read statistics Supplemental Table S2. Orthogroups

Supplemental Table S3. Overrepresented GO terms in *I. aldrichi* singleton genes Supplemental

Table S4. Overrepresented GO terms for genes in *I. aldrichi* species-specific families

Supplemental Table S5. Overrepresented GO terms in gene families with significant gain/loss rates across the phylogeny

Supplemental Table S6. Overrepresented GO terms for gene families expanding in the *I. aldrichi* lineage

Supplemental Table S7. Overrepresented GO terms for gene families contracting in the *I. aldrichi* lineage

## Acknowledgements

We thank Chris Faulk and Carrie Walls of Decorative Genomics for providing sequencing services. We thank Melissa Schreiner (Colorado State University-Extension, Grand Junction, CO, USA) for the gift of the *I. aldrichi* drawing featured in Figure 2, and Ellie R. Hutchison Cervantes for the gift of the parasitized beetle photo (Figure 1b).

## Funding

The study was funded by support from a USDA, Minnesota Department of Agriculture Specialty Crop Block grant (award: 0092341), and the Minnesota Agricultural Experiment Station, University of Minnesota, St. Paul, MN.

## Conflicts of interest statement

The authors declare no conflict of interest.

## Notes

### Competing Interest Statement

The authors have declared no competing interest.

