## Supplemental File S1 for "The genome of *Istocheta aldrichi* (Diptera: Tachinidae), a parasitoid of the Japanese beetle, *Popillia japonica* (Coleoptera: Scarabaeidae)"

Genome Assembly and Evaluation of the Winsome Fly


### Genome Assembly and Evaluation of the Winsome Fly

#### 2025-10-31

An illustration of the Winsome Fly
(*Istocheta aldrichi*), by Melissa Schreiner (Colorado State
University-Extension, Grand Junction, CO).

### Introduction

This document describes the genome assembly pipeline for the Winsome
Fly (*Istocheta aldrichi*), including raw read correction,
assembly, and quality assessment.

#### Step 1: Prepare Reads

```
# Convert BAM File to Fastq
samtools fastq -@ 6 winsomefly.bam > reads.fastq

# Raw Read Correction with Dorado
/home/installs/dorado/dorado correct \
    /home/winsomefly/reads.fastq \
    > /home/winsomefly/corrected_reads.fasta
```

#### Step 2: Genome Assembly with Hifiasm

```
# Assemble genome from corrected reads
/home/installs/hifiasm/hifiasm \
    -o /home/winsomefly/ISTALD \
    -t 32 \
    /home/winsomefly/corrected_reads.fasta

# Convert primary contigs GFA to FASTA for downstream tools
gfatools gfa2fa \
    /home/winsomefly/ISTALD.p_ctg.gfa \
    > /home/winsomefly/ISTALD.p_ctg.fa

# Completeness assessment with Compleasm
compleasm run \
    -a /home/winsomefly/ISTALD.p_ctg.fa \
    -o /home/winsomefly/compleasm_inital_assembly \
    -t 16 \
    -l diptera \
    -L ~/databases
```

#### Step 3: Scaffolding with ntLink

```
ntLink_rounds run_rounds_gaps \
    target=/home/winsomefly/ISTALD.p_ctg.fa \
    reads=/home/winsomefly/reads.fastq \
    t=5 k =32 w=100 rounds=3
# Expected output: /home/winsomefly/ISTALD_ntlink_scaffolds.fa
```

#### Step 4: Removal of haplotigs and overlaps with Purge\_Dups

```
# Map corrected reads to decontaminated assembly
minimap2 -x map-ont -t 32 \
    /home/winsomefly/ISTALD_ntlink_scaffolds.fa \
    /home/winsomefly/corrected_reads.fasta \
    | pigz > /home/winsomefly/reads_vs_assembly.paf.gz

# Coverage stats and cutoffs
cd /home/winsomefly
pbcstat reads_vs_assembly.paf.gz
calcuts PB.stat > cutoffs

# Split assembly and self-align
split_fa ISTALD_ntlink_scaffolds.fa > split.fa
minimap2 -x asm5 -DP split.fa split.fa \
    | gzip -c - > split.self.paf.gz

# Purge duplicates
purge_dups -2 -T cutoffs -c PB.base.cov split.self.paf.gz > dups.bed

# Extract primary and haplotigs
get_seqs -e -p purged dups.bed ISTALD_ntlink_scaffolds.fa

# Outputs:
#   /home/winsomefly/purged.purged.fa   (primary)
#   /home/winsomefly/purged.hap.fa      (haplotigs)
```

#### Step 5: Decontamination

##### Cross-species decontamination with FCS-GX

```
# Identify contigs due to cross-species contamination with FCS-GX
python3 fcs.py screen genome --fasta \
    /home/winsomefly/purged.purged.fa \
    --out-dir /home/winsomefly/fcs_gx \
    --gx-db /home/databases/FCS-GX/gxdb \
    --tax-id 2500616

# Create contaminated contigs list
grep -Eo "ptg[0-9]{6}[a-z]_1" *.txt > contam_contigs.list
wc contam_contigs.list # confirm expected number of contaminated contigs

# Create list of all contigs
grep ">" purged.purged.fa > ISTALD_contigs_all.list
grep ">" purged.purged.fa | wc -l # count original contigs
grep ">" ISTALD_contigs_all.list | wc -l # confirm same number of contigs
sed -i 's/>//g' ISTALD_contigs_all.list # remove >, -i is in place editing

# Generate list of contigs to keep: remove contamination list from all contigs list
grep -v -f contam_contigs.list ISTALD_contigs_all.list > ISTALD_contigs_keep.list # inverse selection in grep

# Check contig counts (new lines) are as expected
wc ISTALD_contigs_keep.list # confirm contaminated contig counts were removed

# Keep only contigs in keep list in the assembly
perl -ne 'if(/^>(\S+)/){$c=$i{$1}}$c?print:chomp;$i{$_}=1 if @ARGV' ISTALD_contigs_keep.list \
  ./purged.purged.fa > ISTALD_assembly.fa
grep ">" ISTALD_assembly.fa | wc -l # confirm final contig count in decontaminated assembly file
```

##### Adaptor and vector decontamination with FCS-adaptor

```
# step 1. screen genome for adapters
./run_fcsadaptor.sh \
--fasta-input ./ISTALD_assembly.fa \
--output-dir FCS_adaptor \
--euk --container-engine singularity --image ./fcs-adaptor.sif

# step 2.  clean genome
cat ./ISTALD_assembly.fa | python3 ./FCS-GX/fcs.py clean genome \
--action-report ./FCS_adaptor/fcs_adaptor_report.txt \
--output ISTALD_assembly_clean.fasta \
--contam-fasta-out ISTALD_contam.fasta
```

#### Step 6: Assembly Assessment

```
# Completeness assessment with Compleasm
compleasm run \
    -a /home/winsomefly/ISTALD_assembly.fa \
    -o /home/winsomefly/compleasm \
    -t 16 \
    -l diptera \
    -L ~/databases


# Evaluation with blobtools
# Map reads to assembly to get alignments, reads after reference genome
minimap2 \
  -t 24 \
  -ax map-ont \
  --secondary=no \
  /home/winsomefly/ISTALD_assembly.fa \
  /home/winsomefly/corrected_reads.fasta \
  -o map.bam

# Sort the mapped reads
samtools sort \
  -@ 24 \
  -o sort.bam \
  map.bam

# Index the sorted BAM file
samtools index sort.bam

# Create a BlobTools database for the primary assembly without BLAST files
blobtools create \
  -i /home/winsomefly/ISTALD_assembly.fa \
  -b ./sort.bam \
  -o ISTALD_blobtools
  --names ./data/names.dmp \
  --nodes ./data/nodes.dmp

# Visualize the results for the primary assembly
blobtools view -i ISTALD_blobtools.blobDB.json
blobtools plot -i ISTALD_blobtools.blobDB.json


# K-merâbased quality assessment with Merqury
# Determine optimal k if needed:
best_k.sh 830134368 0.001

# Create K-mer database from raw reads
meryl k=19 count /home/winsomefly/reads.fastq output /home/winsomefly/read.meryl

# Merqury k-mer assessment of assembly
merqury.sh /home/winsomefly/read.meryl \
    /home/winsomefly/ISTALD_assembly.fa
```

#### Step 7: Generate AGP File and Separate Scaffolds into Contigs with FASTA2AGP

```
java -jar ./FASTA2AGP/src/FASTA2AGP.jar \
  -i ISTALD_assembly_clean.fa \
  -o ISTALD_contig_assembly.fa \
  -a ISTALD_contig_assembly.agp \
  -n 9
```

#### Step 8: Repeat library construction with RepeatModeler

```
BuildDatabase -name primary_db /home/winsomefly/ISTALD_contig_assembly.fa
RepeatModeler -database primary_db -pa 30 -LTRStruct
```

#### Step 9: Repeat masking with RepeatMasker

```
RepeatMasker -lib /home/winsomefly/WF-families.fa \
    -pa 16 \
    -s -xsmall \
    /home/winsomefly/ISTALD_contig_assembly.fa

mv ISTALD_contig_assembly.fa.masked ISTALD_contig_assembly.masked.fa
```

#### Step 10: Structural Protein Annotation with BRAKER

```
singularity exec braker3.sif braker.pl --genome=ISTALD_contig_assembly.masked.fa \
--species=ISTALD \
--prot_seq=Arthropoda.fa \
--AUGUSTUS_CONFIG_PATH=/home/braker/Augustus/config \
--gff3 --threads=32
```

#### Step 11: Check for overlapping gene annotations with AGAT

```
singularity run agat.sif
agat_sp_fix_features_locations_duplicated.pl \
--gff braker.gff3 -o braker.fixoverlaps.gff3
```

#### Step 12: Functional Protein Annotation

##### InterProScan

Functional analysis and database references search with
InterProScan.

```
# remove * from input braker.aa file
cp /home/braker/braker.aa /home/interproscan/braker.aa.fasta # work on a copy with correct filetype extension
sed -i "s/\*//g" braker.aa.fasta # remove all * characters
grep "*" braker.aa.fasta | wc # confirm: should be 0

# run interproscan
./interproscan.sh -i ISTALD_contig_assembly.masked.fa \
-b /home/winsomefly/interproscan \
--goterms \
-t p \
-cpu 18 \
-f TSV, GFF3

# where 
# -b /path/to/output
# --goterms flag to get GO terms
# -t p (as opposed to: n) specifies protein sequences instead of nucleotide
# -cpu indicates 18 cores used for processing, leaving 2 extra available
# -f TSV, GFF3, indicates both gff3 and and TSV file format outputs

# post-processing
# remove everything after '##FASTA' in output gff3 file, this is all protein info
awk '/##FASTA/ {exit} {print}' *.gff3 > temp.gff
# remove header info from gff, by deleting the 1st through 4th line
sed '1,4d' temp.gff > IAL_interpro.gff
rm edit.gff
```

##### eggNOG-mapper

Additionally, run a secondary gene function search with
eggNOG-mapper.

```
# step 1: search for proteins with diamond in VERY-sensitive mode
emapper.py -i /home/winsomefly/interproscan/braker.aa.fasta \
-m diamond \
--sensmode very-sensitive \
--no_annot \
--go_evidence experimental \
-o eggnog \
--excel \
--cpu 0 

# step 2: annotate by loading annotation DB entirely into memory with --dbmem (44 GB free memory)
emapper.py -m no_search \
--cpu 0 \
--annotate_hits_table IAL.emapper.seed_orthologs \
-o IAL.emapper_annot \
--excel \
--decorate_gff yes \
--dbmem

# where
# -cpu 0 indicates to run with all available CPUs
# -o indicates output file base name prefix 
# --decorate_gff yes: a new GFF will be created, and decorated with hits and/or annotations
# --excel also output annotations in an .xlsx format
# --go_evidence experimental or non-electronic indicates which GO terms to include (default is non-electronically curated)
# -dbmem indicates high GB RAM available, >44 can increase annotation speed considerably
```

##### Append protein functions to structural protein GFF

Clean and combine functional annotation information from both
programs: InterProScan and eggNOG-mapper.

```
# Define your input and output files
interproFile="IAL_interpro.gff"
eggFile="IAL_annot.emapper.decorated.gff"
outFile="~/home/IAL_combined_functional_annotations.csv"

# Set Up R environment
library('stringr')
library('dplyr') 
setwd("~/home/winsomefly")

# import gff files into R and remove headers
interpro <- read.delim(interproFile, sep="\t", header = FALSE)
interpro2 <- interpro %>% 
  filter(V3 != "polypeptide") %>% # remove rows indicating polypeptide itself
  filter(!(grepl('#', V1))) # grep : search for substring match in V1 column
interpro <- interpro2
rm(interpro2)

eggnog <- read.delim(eggFile, sep="\t", header = FALSE)
eggnog2 <- eggnog %>% 
  filter(!(grepl('#', V1))) # remove headers
eggnog <- eggnog2
rm(eggnog2)

# Extract database references from interpro
interpro$Dbxref <- str_remove(interpro$V9, ".*Dbxref=") # removes everything until keyword
interpro$V10 <- str_remove_all(interpro$Dbxref, '"') # removes all quotation marks

interpro$V11 <- str_extract(interpro$V9, "(?<=Ontology_term=).*?(?=\\;)") # extract Go Terms
interpro$V12 <- str_remove_all(interpro$V11, '"') # removes all quotation marks

interpro2 <- interpro %>% 
  mutate(V10 = if_else(str_detect(V10, "InterPro:"), V10, NA_character_)) %>% # if no match make value NA
  select(!(Dbxref|V11)) # remove temp columns

# Collect unique InterPro terms and GoTerms for each gene

# summarize into one row per gene
interpro_combined <- interpro2 %>% 
  filter(!is.na(V10)) %>% 
  group_by(V1) %>% 
  summarize(DbxRef = paste(V10, collapse = ","), GoTerms = paste(V12, collapse = ","))

# removes NA text string, replaces true NAs to NA value
interpro_combined$GoTerms <- str_remove_all(interpro_combined$GoTerms, 'NA,')
interpro_combined$GoTerms <- str_remove_all(interpro_combined$GoTerms, ',NA') 
interpro_combined$GoTerms[interpro_combined$GoTerms == "NA"] <- NA

# remove repeated database reference terms
remove_repeats <- function(x) {
  Terms <- unlist(strsplit(x, ","))
  uniqueList <- unique(Terms)
  paste(uniqueList, collapse = ",")
}
interpro_combined$uniqueGoTerms <- sapply(interpro_combined$GoTerms, remove_repeats)
interpro_combined$uniqueDbxref <- sapply(interpro_combined$DbxRef, remove_repeats)

# keep only needed data from InterProScan
interpro <- interpro_combined %>% 
  select(V1,uniqueGoTerms,uniqueDbxref)
rm(interpro_combined)
interpro$uniqueGoTerms[interpro$uniqueGoTerms == "NA"] <- NA

# Extract description text from eggNOG mapper
eggnog$V10 <- str_extract(eggnog$V9, "(?<=em_desc=).*?(?=\\;)")
eggnog$V11 <- str_remove(eggnog$V9, ".*em_GOs=")
eggnog2 <- eggnog %>% 
  mutate(V11 = if_else(str_detect(V11, "GO:"), V11, NA_character_)) %>% # if not match make value NA
  select(!(V9))


# Filter join
# Collect all GO terms for shared annotations in one dataframe!
rm(eggnog)
df <- merge(x = eggnog2, y = interpro, by = "V1", all.x = TRUE)

# combine GoTerms to one column
df <- df %>% 
  mutate(V11 = case_when(
    !is.na(V11) & !is.na(uniqueGoTerms) ~ paste(V11, uniqueGoTerms, sep = "-"),
    !is.na(V11) ~ V11,
    !is.na(uniqueGoTerms) ~ uniqueGoTerms,
    TRUE ~ NA_character_
  ))
# remove repeats
df$GoTerms <- sapply(df$V11, remove_repeats)
df$GoTerms[df$GoTerms == "NA"] <- NA

# Paste components into new final attributes column
df$attributes <- paste0("ID=", df$V1, ";desc=", df$V10, ";Dbxref=", df$uniqueDbxref, ";GO_terms=", df$GoTerms)

# select columns to keep 
df2 <- df %>% 
  select(1:8,14)
rm(df)

# add in annotations from InterPro that eggNOG did not catch
# select annotation source with highest score from interproscan
interproWinner <- interpro2 %>%
  group_by(V1) %>%
  slice_max(V6, n = 1, with_ties = FALSE)

# find all transcripts that are in interproWinner but NOT eggnog + interpro supplemented gff
interproOnly <- anti_join(interproWinner, df2, by = "V1")

# extract signature descriptions
interproOnly$sigdesc <- str_extract(interproOnly$V9, "(?<=signature_desc=).*?(?=\\;)") 

# Paste components into new final attributes column
interproOnly$attributes <- paste0("ID=", interproOnly$V1, ";desc=", interproOnly$sigdesc, ";Dbxref=", interproOnly$V10, ";GO_terms=", interproOnly$V12)

# remove extra temp rows
interproToAppend <- interproOnly %>% 
  select(!(9:12)) # remove temp columns

# append functional annotations from InterProScan to create aggregated GFF file with functional annotations
gff <- rbind(df2, interproToAppend)

# format change from interpro output: protein_match -> CDS
gff$V3 <- gsub("protein_match", "CDS", gff$V3)

# export joined functional annotations to csv
write.csv(gff, outFile, row.names = FALSE)
```

Append the functional annotations to the structural annotation and
output one GFF file.

```
##### Append Functional Annotations to Structural GFF ####

noDescEGG <- gff %>% 
  filter(grepl("desc=None;Dbxref=NA;GO_terms=NA", attributes)) # grepl : search for substring match in attributes column

noDescINT <- gff %>% 
  filter(grepl("desc=NA;Dbxref=NA;GO_terms=NA", attributes))

gff2 <- gff %>% 
  mutate(attributes = gsub("desc=None", "desc=NA", attributes)) # change empty eggNOG desc to NA

gff2 <- gff2 %>% 
  mutate(attributes = if_else(str_detect(attributes, "desc=NA;Dbxref=NA;GO_terms=NA"), NA_character_, attributes)) # if match (no func. anno) then make value NA, else keep attributes 

# clean up disordered region flags, as these are not strictly functions
gff2 <- gff2 %>% 
  mutate(attributes = if_else(str_detect(attributes, "desc=consensus disorder prediction;Dbxref=NA;GO_terms=NA"), NA_character_, attributes))

gff2 <- gff2 %>% 
  mutate(attributes = gsub("ID=", "Name=", attributes))

# Add functional gene annotations to final GFF
strucGFF <- read.delim("braker_overlap_fixed_20251013.gff3", sep="\t", header = FALSE)

strucGFF$seqID <- strucGFF$V1
strucGFF$V1 <- str_extract(strucGFF$V9, "(?<=ID=).*?(?=\\;)") # extract gene ID

df <- merge(x = strucGFF, y = gff2, by = "V1", all.x = TRUE)

# remove extraneous terms eg ";Go_terms=NA"
df <- df %>% 
  mutate(attributes = gsub(";GO_terms=NA", "", attributes))

df <- df %>%
  mutate(attributes = gsub(";Dbxref=NA", "", attributes))

# Paste components into new final attributes column
df$attributes2 <- if_else(!is.na(df$attributes), paste0(df$V9, df$attributes), NA)
df$V9 <- if_else(!is.na(df$attributes2), paste0(df$attributes2), paste0(df$V9))

df$V1 <- df$seqID

# select columns to keep 
df2 <- df %>% 
  select(1:9)
rm(df)

# rename column names for a gff file
colnames(df2) <- NULL
# colnames(gff) <- c("seqID","source","type","start","end","score","strand","phase","attributes")

# export to csv
write.csv(df2, "~/Downloads/IAL_GFF_strucAndFunct.csv", row.names = FALSE)

# export to .gff
write.table(
  df2,
  file = "IAL_annotated.gff3", # Name of the output file
  sep = "\t",          # Specify tab as the separator
  row.names = FALSE,   # Do not include row names in the file
  col.names = FALSE,    # Include column names (header) in the file
  quote = FALSE        # Do not quote character strings
)
```

#### Step 13: Rename genes with species prefix in AGAT

```
singularity run agat.sif 
agat_sp_manage_IDs.pl \
--gff IAL_annotated.gff3 \
--prefix IAL --ensembl TRUE --type_dependent -o IAL_ensembl.gff3
exit

# remove extraneous leading zeros 
cp IAL_ensembl.gff3 IAL_renamed.gff3
sed -i 's/IALG000000/IAL_/g' IAL_renamed.gff3

# generate protein sequences from renamed genes with AGAT (.aa file)
agat_sp_extract_sequences.pl \
-g IAL_renamed.gff3 \
-f ISTALD_contig_assembly.masked.fa \
-p \
-o IAL_protein.aa
```

#### Step 13: Prepare Annotated Assembly for Submission to GenBank

```
./table2asn -M n -J -c w -euk \
  -t ./template.sbt \
  -j "[organism=Istocheta aldrichi]" \
  -i ./ISTALD_contig_assembly.masked.fa \
  -f ./IAL_renamed.gff3 \
  -o IAL.sqn \
  -locus-tag-prefix IAL \
  -usemt many \
  -Z -V b
```

#### Step 14: Compute Statistics on Final Assembly

```
# Table 2. Istocheta aldrichi genome assembly and curation statistics
FASTA=ISTALD_contig_assembly.masked.fa 
SCAFFOLDS=ISTALD_assembly_clean.fa
/home/installs/seqkit stats -a ${FASTA} > contigs.stats # -a flag gives L50 ("N50_num")
/home/installs/seqkit stats -a ${SCAFFOLDS} > scaffolds.stats


# Table 3. Interspersed repeats in I. aldrichi 
# generated during repeatmodler run (see Step 8)
cd home/winsomefly/repeatmodeler/
cat *.tbl


# Table 4. Istocheta aldrichi genome annotation metrics with GAG
python /home/installs/GAG/gag.py \
--fasta /home/winsomefly/ISTALD_contig_assembly.masked.fa \
--gff IAL_renamed.gff3 \
--out GAG_IAL
```

### Mitochondrial assembly

#### Step 1: Fetch mitochondrial reference & Assemble mitochondrial genome with MitoHiFi

```
mkdir -p ~/scripts/outputs/mitohifi_results/references

findMitoReference.py \
  --species "Istocheta aldrichi" \
  --outfolder ~/scripts/outputs/mitohifi_results/references \
  --min_length 14000

mitohifi.py \
  -r /home/winsomefly/corrected_reads.fasta \
  -f ~/scripts/outputs/mitohifi_results/references/OR526344.2.fasta \
  -g ~/scripts/outputs/mitohifi_results/references/OR526344.2.gb \
  -t 16
```

#### Step 2: Circularization start correction (Circlator)

```
circlator fixstart \
  ~/scripts/outputs/mitohifi_results/final_mitogenome.fasta \
  ~/scripts/outputs/mitohifi_results/circularized.fasta
```

### Gene Family Size Changes Analysis

To create Figure 3, the following code was used to characterize
GoTerms for each gene family.

```
#!/usr/bin/perl
use strict;
use warnings;

my @inputb2g = glob("/home/winsomefly/proteins/*.goList.txt");
##For each species, you need a tab delimited file in which go terms are listed for each gene as such:
#e.g.:
#IALM00000000003    0004190,0006508,0003676,0008270

my $inputortho = "/home/winsomefly/proteins/OrthoFinder/Results/Orthogroups/Orthogroups4GOassign.txt";
#Provide a tab delimited file in which the name of the orthogroup/gene family is in the first column, and the list of genes in the second column, space separated 
#e.g.:
#OG0024884: TRIPEN_g21662.t1 TRIPEN_g21739.t1

my $output = "/home/winsomefly/proteins/OrthoFinder/Results/Orthogroups/GO_terms_by_gene_family.txt";
my %list;

foreach my $file (@inputb2g) {    ##############goes through each of the go list txt files
    $file =~ /proteins\/(\w+).goList/; ########## grabs the species name that begins the name of each B2G txt file ex., /Blast2GP/APME_b2g.... =APME
    my $species = $1;             ######### assigns the previous line to $species
    open FILE, "$file";     ######### opens the file that we just grabbed the name from
    while (my $line = <FILE>) { ######## go line by line
        my ($gene, $gos) = split /\t/, $line;   ########## split the line on tab. First column = gene, second = list of go terms for that gene
        $list{$gene} = $gos;              ############ now put the gene name and associated go terms into a hash, where gene name = key & list of GOs = value
        }
    close FILE;             ####### close file
    }

open ORTHOFILE, "$inputortho";          #####open the file with orthogroup membership information 
open OUTFILE, ">$output";           #####open the output file and tell it we are going to print to it   
while (my $ortholine = <ORTHOFILE>) {       ######### go line by line
    my $count = 0;              ####### for keeping track of the number of genes in each family
    my %gocounts;               ######keep track of the number of times go terms appear within a gene family
    my ($family, $members) = split /:/, $ortholine;   ############split the line on the colon. First column = gene family, second column  = all gene members of the family
    my @familymembers = split /\s/, $members;       ######### split the list of family members and put it into an array
    shift @familymembers;               ######### the first element of the array is a space, shift gets rid of this ###### CHECK THIS 
    foreach my $member (@familymembers) {
        $count++;                   #######for each gene ($member) in @familymembers, increase count by one. To be used later for calculating 40%
        if ($list{$member}) {
            my @gotermsformember = split /,/, $list{$member}; ####### make an array that is all the go terms for the $member in question
            foreach my $goterm (@gotermsformember) {      #####for these go terms for the gene
                if ($gocounts{$goterm}) {       #######see if that go term is already in the hash which is keeping track for the whole gene family  
                    $gocounts{$goterm}++;       ###### if it is already there, increase the count by one
                } else {                #### OR
                    $gocounts{$goterm} = 1;     ##### add it to the hash and set it to one  
                    }
                }
            }
        }

    foreach my $goterm (keys %gocounts) {           ###### new variable $goterm (reusing name, okay b/c previous instance loop was closed), look up the keys of the counts (which is $goterm)
        if ($gocounts{$goterm} / $count >= 0.001) { ##### if the counts of the go term is greater than or equal to  40% of the #of genes in the family...
            print OUTFILE "$family = $goterm\n";    ##### then print it to the outfile in the family = goterm format that is needed for bingo
            }
        }
    }

close ORTHOFILE;
close OUTFILE;
```
